# Sex differences in Variability of Brain Structure Across the Lifespan

**DOI:** 10.1101/842567

**Authors:** Natalie J Forde, Jerrold Jeyachandra, Michael Joseph, Grace R Jacobs, Erin Dickie, Theodore D Satterthwaite, Russell T Shinohara, Stephanie H Ameis, Aristotle N Voineskos

**Author notes:** **Corresponding author:**, Centre for Addiction and Mental Health, 250 College St, M5T 1R8, Toronto ON, Canada.

## Abstract

Several brain disorders exhibit sex differences in onset, presentation, and prevalence. Increased understanding of the neurobiology of sex-based differences across the lifespan can provide insight into potential disease risk and protective mechanisms. We focused on sex-related differences in variability, which may be indicative of both disease vulnerability and resilience. In n=3,069 participants, from 8-95 years of age, we first analyzed the variance ratio in females vs. males of cortical surface area and global and subcortical volumes for discrete brain regions, and found widespread greater variability in males. In contrast, variance in cortical thickness was similar for males and females. Multivariate analysis that accounts for structural covariance supported variance ratio findings. Findings were present from early life and stable with age. We then examined variability among brain regions by sex. We found significant age-by-sex interactions across neuroimaging metrics, whereby in very early life males had reduced among-region variability compared to females, while in very late life this was reversed. Overall, our findings of greater regional variability but less among-region variability in males in early life may aid our understanding of sex-based risk for neurodevelopmental disorders. In contrast, our findings in late life may provide a potential sex-based risk mechanism for dementia.

## Introduction

Sex differences in onset, presentation, and prevalence are common in many brain disorders (1). For instance, the prevalence of autism spectrum disorders (ASD; (2)) is increased 4-5 fold in males compared to females. Schizophrenia prevalence is also increased in males; age of onset and illness presentation also varies between males and females (3, 4). In contrast, depression and Alzheimer’s Dementia are more prevalent in females compared to males (5–7). A better understanding of sex-related differences in healthy brain architecture at different phases of the lifespan may help identify risk factors and protective mechanisms for psychiatric and neurologic disorders.

To date, studies comparing the brain in males and females have focused primarily on identifying differences, on average, between groups (e.g. (8–10)). Findings from such studies have often generated controversy (11, 12) and frequently fail to fully address the complexity of the topic (13). For instance, when taking group-averages, males tend to have larger total brain volumes than females; however the majority of regional differences reported can be attributed to the difference in total brain volume (14, 15). Others have shown differences in functional network organization and cerebral blood flow between males and females (9), which are likely unrelated to brain volume. However, a focus on group differences between the sexes may underplay the importance of the considerable heterogeneity and overlap between and among the sexes (13).

There is increasing recognition of the importance of variability in brain structure and function in both health and disease (15–17). Recent work in older adults shows that males have greater variability than females across indices (surface area and volumes) of brain structure even after accounting for total brain volume (15). This study provided further evidence that the majority of differences, on average, between males and females can be attributed to variance in total brain volume. A second study, using a developmental cohort (n=1,234), similarly found greater variance in males compared to females in volumes of multiple subcortical structures (17). These consistent findings at opposing ends of the lifespan are of particular consequence when considering the differences in prevalence between the sexes in certain brain disorders (e.g. neurodevelopmental disorders), and the timing of their onset and course (2). However, brain disorders are increasingly understood as ones where relationships between or among brain regions are disrupted. Examining relationships among regions (18) has illuminated our understanding of brain organization across the lifespan, and in brain disorders (19, 20). However, the variability of such relationships among regions, is to our knowledge, not known in males or females.

Here we analyze data in over 3,000 participants from three large, high-quality, open-source datasets to investigate structural variability across the lifespan. Notably these datasets are independent of the datasets used previously to investigate brain structure variability (15, 17). The current study can offer insights into periods of rapid reorganization and development in the brain from: childhood through adolescence (Philadelphia Neurodevelopmental Cohort [PNC] (21)), a relatively stable period during young adulthood following completion of the majority of developmental processes (22, 23) (Human Connectome Project [HCP] (24)), and the re-emergence of dynamic change that occurs in late life (25) as part of the aging process (Open Access Series of Imaging Studies [OASIS-3] (26–28)).

In the present study we aim to comprehensively examine structural variability across the lifespan. Our first aim was to examine sex-based variability by region and measurement type (surface area, cortical thickness, subcortical volume). Based on recent literature, (15, 17), we hypothesized that greater variability would be present in surface area and volume measures in males compared to females. These are measures of brain structure under strong genetic control and are largely determined early in development, unlike cortical thickness which is under more considerable environmental control (29). Our second aim was to investigate sex-based variability in relationships among brain regions. If higher in one sex compared to the other, this is indicative of an elevated degree of variability in the relationships of brain regions to each other within that sex. As opposed to network approaches that focus on individual connections, nodes or subnetworks, the current approach considers the overall pattern of relationships between all regions within an individual. In this way it complements standard variability analysis that considers regions independently of each other rather than the relationships between them. We hypothesized that such variability would be age dependent and align with differential risk for complex brain disorders at either end of the lifespan, consistent with age dependent network reorganization (30–32) and altered relationships among structures in these disorders.

## Results

To assess sex differences in variance we analysed total brain volumes, subcortical volumes, regional surface area, and regional cortical thickness measures from the PNC (8-21 years) dataset (n=1,347), the HCP Young Adult (22-37 years) S1200 dataset (N=1,032) and the OASIS-3 (42-95 years) dataset (n=609). See Materials and Methods for details.

### Variance Ratio Across Measures and Regions

For the first aim, we regressed age from each metric to focus on sex differences and compared the variance ratio between sexes for each region. Analyses were also repeated with total brain volume additionally regressed out to determine if differences in total brain volume between the sexes accounted for regional findings.

#### Global Volume

Across all datasets we found total brain volume, cerebral grey matter volume, and cerebral white matter volume were more variable in males compared to females (VR>1, *p*<0.1; Table 1, Figure 1). In the young adult (HCP) and late-life (OASIS-3) datasets cerebellar white matter volume showed a similar pattern of higher variance in males compared to females, but this was not found in the child and youth (PNC) dataset. Cerebellar grey matter volume showed a similar but non-significant pattern of increased variability in males compared to females across all datasets. Accounting for total brain volume did not change any of these results.

**Table 1.** Variance in Global Volumes.

|  |  | PNC |  | HCP |  | OASIS-3 |  |
| --- | --- | --- | --- | --- | --- | --- | --- |
|  |  | VR | q | VR | q | VR | q |
| **TBV** | raw | 1.20 | 0.09 | 1.36 | **0.002** | 1.38 | **0.001** |
| **Cerebral GM** | raw | 1.18 | 0.09 | 1.31 | **0.003** | 1.32 | **0.003** |
|  | corrected | 1.22 | **0.02** | 1.36 | **8.4E-04** | 1.41 | **2.5E-04** |
| **Cerebral WM** | raw | 1.25 | **0.02** | 1.42 | **9.7E-05** | 1.45 | **1.4E-04** |
|  | corrected | 1.25 | **0.02** | 1.32 | **0.002** | 1.37 | **6.8E-04** |
| **Cerebellar GM** | raw | 1.13 | 0.19 | 1.15 | 0.13 | 1.07 | 0.54 |
|  | corrected | 1.20 | **0.03** | 1.14 | 0.15 | 1.10 | 0.36 |
| **Cerebellar WM** | raw | 0.97 | 0.89 | 1.49 | **1.4E-05** | 1.39 | **4.6E-04** |
|  | corrected | 0.99 | 1 | 1.64 | **2.4E-08** | 1.58 | **1.1E-06** |

Results from analyses of sex differences for global brain volumes are presented from the Philadelphia Neurodevelopmental Cohort (PNC), Human Connectome Project (HCP) and Open Access Series of Imaging Studies (OASIS-3). Variance ratios (VR) were calculated from F-tests, VR>1 indicates males > females and VR<1 indicates females > males in variance. 'Raw' results represent findings from volumes corrected for age (age effects regressed out). Corresponding 'corrected' results were generated from volumes corrected for total brain volume (TBV) as well as age. GM - grey matter, WM - white matter, *q* - False Discovery Rate (FDR) corrected *p*-value.

**Figure 1.**
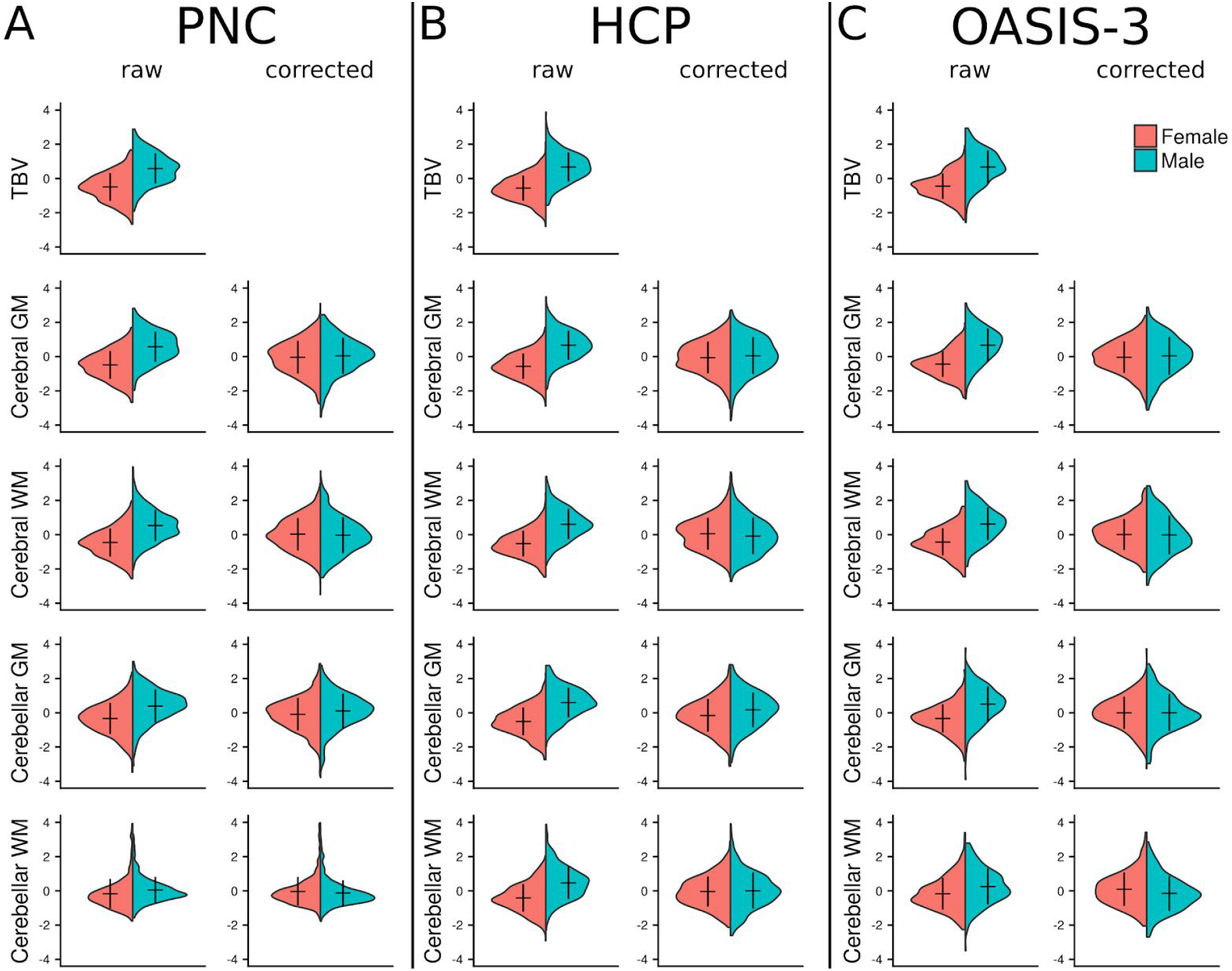
Sex Differences on Global Volume. Global brain volume measures from three independent datasets are shown; (A) The Philadelphia Neurodevelopmental Cohort (PNC), (B) The Human Connectome Project (HCP) and (C) The Open Access Series of Imaging Studies (OASIS-3). ‘Raw’ data represent volumes corrected for age (following regression of age from the model). Corresponding ‘corrected’ figures show volume data corrected for total brain volume (TBV) as well as age. Mean and standard deviation of the data are represented by the horizontal and vertical lines, respectively. See Table 2 for full statistical results. GM - grey matter, TBV - total brain volume, WM - white matter.

#### Subcortical Volume

Across all three datasets, we found significant variance ratio differences by sex in subcortical brain volumes, with males exhibiting greater variance compared to females across most regions, while there were no regions in which females were greater than males (Table 2, Figure 2 top row). The pattern of findings was similar following correction for total brain volume. There were, however, some differences in findings across datasets. For instance, variance ratios were not significantly different by sex in hippocampus and nucleus accumbens volumes in the child and youth PNC dataset, although we did find significance in the young-adult HCP (left hippocampus and bilateral nucleus accumbens) and later-life OASIS-3 (bilateral hippocampus and nucleus accumbens) datasets, with males showing greater variance compared to females. Putamen and pallidum volumes did show significant variance ratios by sex in the PNC and OASIS-3 datasets (males > females), while there were no significant differences in the HCP data.

**Table 2.** Variance in subcortical volumes.

|  |  |  | PNC |  | HCP |  | OASIS-3 |  |
| --- | --- | --- | --- | --- | --- | --- | --- | --- |
|  |  |  | VR | q | VR | q | VR | q |
| **Thalamus** | left | raw | 1.14 | 0.15 | 1.27 | **0.01** | 1.22 | 0.09 |
|  |  | corrected | 1.41 | **1.3E-04** | 1.17 | 0.09 | 1.09 | 0.44 |
|  | right | raw | 1.28 | **0.02** | 1.35 | **0.002** | 1.84 | **9.4E-07** |
|  |  | corrected | 1.32 | **0.003** | 1.26 | **0.03** | 1.52 | **8.6E-04** |
| **Caudate** | left | raw | 1.26 | **0.02** | 1.25 | **0.02** | 1.69 | **2.5E-05** |
|  |  | corrected | 1.25 | **0.01** | 1.11 | 0.30 | 1.64 | **1.4E-04** |
|  | right | raw | 1.18 | 0.07 | 1.31 | **0.005** | 1.58 | **2.2E-04** |
|  |  | corrected | 1.17 | 0.07 | 1.20 | 0.07 | 1.53 | **8.1E-04** |
| **Putamen** | left | raw | 1.20 | 0.05 | 1.09 | 0.34 | 1.59 | **2.2E-04** |
|  |  | corrected | 1.14 | 0.14 | 1.09 | 0.38 | 1.59 | **3.2E-04** |
|  | right | raw | 1.21 | 0.05 | 1.16 | 0.11 | 1.86 | **9.4E-07** |
|  |  | corrected | 1.21 | **0.02** | 1.11 | 0.30 | 1.74 | **2.3E-05** |
| **Pallidum** | left | raw | 1.18 | 0.07 | 1.05 | 0.60 | 1.36 | **0.01** |
|  |  | corrected | 1.26 | **0.01** | 0.99 | 0.89 | 1.33 | **0.03** |
|  | right | raw | 1.20 | 0.05 | 1.09 | 0.34 | 1.35 | **0.01** |
|  |  | corrected | 1.25 | **0.01** | 1.06 | 0.56 | 1.24 | 0.07 |
| **Hippocampus** | left | raw | 0.98 | 0.80 | 1.55 | **3.9E-06** | 1.37 | **0.009** |
|  |  | corrected | 0.96 | 0.66 | 1.68 | **1.5E-08** | 1.25 | 0.06 |
|  | right | raw | 1.03 | 0.78 | 1.17 | 0.10 | 1.53 | **5.1E-04** |
|  |  | corrected | 1.03 | 0.66 | 1.22 | 0.05 | 1.41 | **0.008** |
| **Amygdala** | left | raw | 1.11 | 0.27 | 1.39 | **7.0E-04** | 1.56 | **2.7E-04** |
|  |  | corrected | 1.23 | **0.02** | 1.31 | **0.01** | 1.30 | **0.04** |
|  | right | raw | 1.12 | 0.21 | 1.24 | **0.02** | 1.44 | **0.003** |
|  |  | corrected | 1.19 | 0.05 | 1.17 | 0.09 | 1.32 | **0.03** |
| **Nucleus**<br>**Accumbens** | left | raw | 1.10 | 0.29 | 1.24 | **0.02** | 1.27 | **0.04** |
|  |  | corrected | 1.12 | 0.17 | 1.24 | **0.04** | 1.26 | 0.06 |
|  | right | raw | 1.07 | 0.48 | 1.25 | **0.02** | 1.41 | **0.004** |
|  |  | corrected | 1.07 | 0.45 | 1.17 | 0.09 | 1.36 | **0.02** |

Results from analyses of sex differences in subcortical volumes are presented from the Philadelphia Neurodevelopmental Cohort (PNC), Human Connectome Project (HCP) and Open Access Series of Imaging Studies (OASIS-3). Variance ratios (VR) were calculated from F-tests, VR>1 indicates males > females and VR<1 indicates females > males in variance. 'Raw' results represent findings from volumes corrected for age (following regression of age from the model). Corresponding 'corrected' results were generated from volumes corrected for total brain volume (TBV) as well as age. *q* - False Discovery Rate (FDR) corrected *p*-value

**Figure 2.**
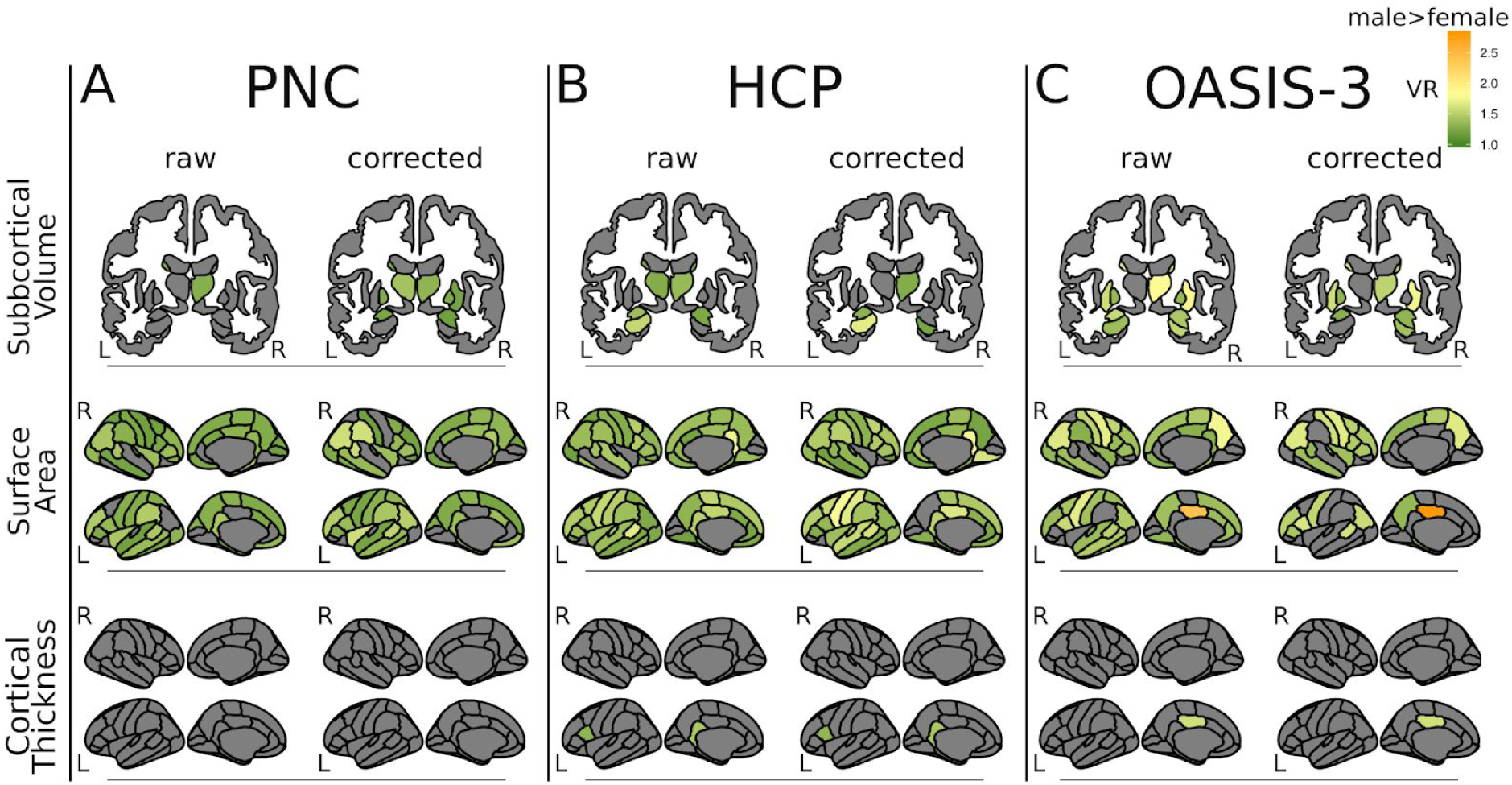
Sex Differences in Variance. Variance ratio (VR) results are mapped onto subcortical structures (top row) or cortical surface (middle and bottom panels) for three independent datasets; (A) The Philadelphia Neurodevelopmental Cohort (PNC), (B) The Human Connectome Project (HCP) and (C) The Open Access Series of Imaging Studies (OASIS-3). Variance Ratios were generated with F-tests comparing metrics of subcortical volume, cortical surface area and cortical thickness between males and females. Only VR’s that met statistical significance are plotted (*q*<0.05). VR>1 (green-yellow-orange) indicates males > females. There were no regions where females had greater variance than males (VR<1). ‘Raw’ figures show results from analyses that used metrics corrected for age (following regression of age from the model). Corresponding ‘corrected’ figures show results from analyses that used metrics corrected for total brain volume (TBV) as well as age. Note: Nucleus Accumbens is not included in the figure but does show significant differences in variability between males and females (males higher) in the HCP and OASIS-3 sample (see Table 3). L - left, R- right.

#### Surface Area

Significant variance ratios by sex in surface area were found across the great majority of cortical regions, whereby males demonstrated greater variance compared to females across all datasets (Figure 2, middle 2 rows), while there were no regions in which females were greater than males. In the child and youth PNC dataset, we found significant variance ratios that in 52/68 regions (VR 1.17-1.56, *q*<0.05 [57/68, *q*<0.1]), all 52 of which males > females. In the HCP young adult dataset, males had higher variance in 59/68 regions (VR 1.19-2.14, *q*<0.05 [61/68, *q*<0.1]). The OASIS-3 later-life dataset similarly displayed higher variance in males compared to females in 44/68 regions (VR 1.29-2.40, *q*<0.05 [49/68, *q*<0.1]). Findings in all datasets were similar following correction for total brain volume (PNC, 51/68 regions, VR=1.19-1.61, *q*<0.05; HCP, 55/68 regions, VR=1.19-2.15, *q*<0.05; OASIS-3, 27/68 regions, VR=1.32-2.80, *q*<0.05).

#### Cortical Thickness

Variance in cortical thickness was similar between males and females across all datasets (Figure 2, bottom 2 rows), with variance ratios significant in a very small number of regions. Only portions of the left cingulate gyrus showed higher variance in males compared to females in adults in the young adult HCP dataset (VR=1.44, *q*=0.001) and in the OASIS-3 later-life dataset (VR=1.59, *q*=0.004). Additionally the left pars opercularis was significantly more variable in males compared to females in the HCP dataset (VR=1.32, *q*=0.04). These effects remained consistent following correction for total brain volume.

### Mahalanobis Distance

The Mahalanobis distance is a single measure per participant, which incorporates all metrics in the model, and represents a participant’s difference from the group average. Consistent with variance ratio analyses, Mahalanobis distance is used here as a measure of variance within the population. However, it provides a more statistically robust approach by accounting for the covariance between regions. We grouped our metrics by type: global volumes (n=4), subcortical volumes (n=14), surface area (n=68), and cortical thickness (n=68) and calculated Mahalanobis distance for each participant in relation to their group average. Males demonstrated greater Mahalanobis distance in global volume, subcortical volume and surface area compared to females across all datasets (Table 3, Figure 3). In each dataset, surface area showed the largest sex effect, followed by subcortical volumes and then global volumes (S Table 1). There were no significant sex differences in cortical thickness. These results are consistent with the univariate analyses of variance ratio. Analyses with age showed a significant age-by-sex interaction in surface area of the OASIS-3 sample (F=7.18, *q*<0.001), whereby male Mahalanobis distance is greater over the majority of the age distribution but within the oldest age bin (75-80 years) females have greater Mahalanobis distance. No other age-by-sex interactions were found. Significant associations between Mahalanobis distance and age were found across cortical metrics in the OASIS-3 sample; surface area (F=16.23, *q*<0.001) and cortical thickness (F=14.88, *q*<0.001). Cortical thickness Mahalanobis distance followed a non-linear course first slightly increasing with age before declining in later years. In surface area, the younger-old participants (55-60) had a lower Mahalanobis distance, which then increased in later years.

**Table 3.** Mahalanobis Distance.

|  |  | PNC |  | HCP |  | OASIS-3 |  |
| --- | --- | --- | --- | --- | --- | --- | --- |
|  |  | F | q | F | q | F | q |
| **Global** | Sex | 5.42 | **0.02** | 43.26 | **2.3E-10** | 13.83 | **3.3E-04** |
|  | Age | 0.13 | 1.00 | 1.49 | 0.68 | 0.05 | 1.00 |
|  | Age-by-se<br>x | 1.17 | 0.31 | 0.61 | 0.55 | 0.80 | 0.52 |
| **Subcortical Volume** | Sex | 11.63 | **6.7E-04** | 44.40 | **6.5E-11** | 46.63 | **6.5E-11** |
|  | Age | 0.10 | 0.90 | 0.53 | 0.90 | 0.46 | 0.90 |
|  | Age-by-se<br>x | 0.81 | 0.44 | 0.67 | 0.51 | 2.34 | 0.05 |
| **Surface Area** | Sex | 161.02 | **1.2E-34** | 229.86 | **1.4E-46** | 43.89 | **9.4E-11** |
|  | Age | 0.39 | 0.67 | 3.28 | 0.06 | 16.23 | **5.4E-12** |
|  | Age-by-se<br>x | 2.67 | 0.07 | 1.40 | 0.25 | 7.18 | **1.3E-05** |
| **Cortical Thickness** | Sex | 0.85 | 0.36 | 1.41 | 0.35 | 1.77 | 0.35 |
|  | Age | 0.01 | 0.99 | 0.05 | 0.99 | 14.88 | **5.4E-11** |
|  | Age-by-se<br>x | 0.55 | 0.58 | 0.07 | 0.94 | 0.60 | 0.66 |

Mahalanobis distances distributions for three independent datasets; Philadelphia Neurodevelopmental Cohort (PNC), Human Connectome Project (HCP) and Open Access Series of Imaging Studies (OASIS-3) were compared between males and females with a linear model followed by type 2 F-tests. *q* - False Discovery Rate (FDR) corrected *p*-value. All significant sex differences correspond to larger Mahalanobis distance in males compared to females. Mahalanobis distances were calculated for each subject to their group (male or female) average per metric type.

**Figure 3.**
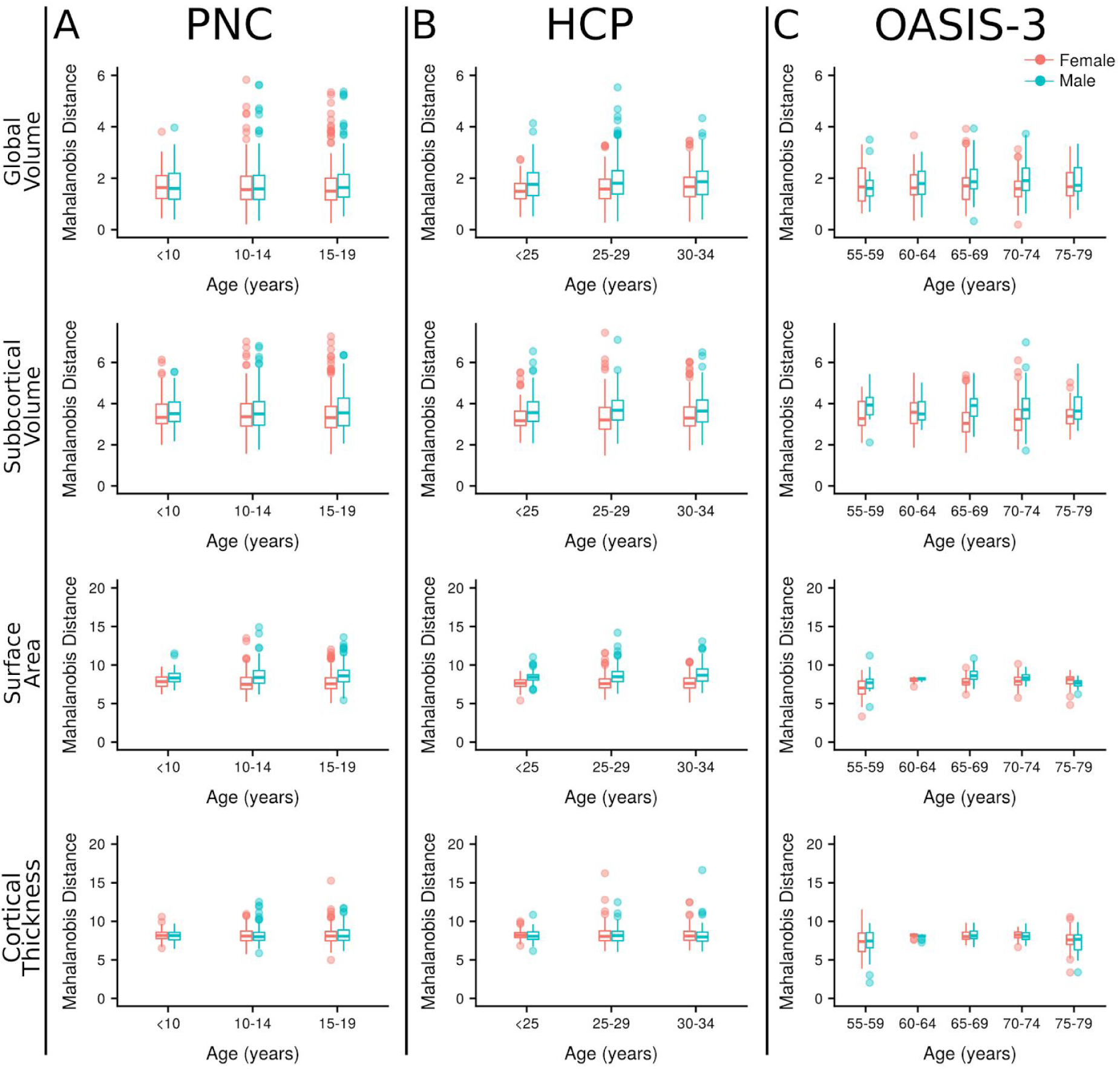
Mahalanobis Distance. Mahalanobis distance plotted by age for subjects from three independent datasets; (A) The Philadelphia Neurodevelopmental Cohort (PNC), (B) The Human Connectome Project (HCP) and (C) The Open Access Series of Imaging Studies (OASIS-3). Mahalanobis distance was calculated for each subject in relation to their group (male or female) average. Metrics included in each analysis were grouped by type; global volumes (n=4), subcortical volumes (n=14), surface area (n=68) and cortical thickness (n=68).

### Cosine Angle Dissimilarity

We then considered the relationship of regions among each other within a subject, and the dissimilarity of these profiles across participants within one sex compared to the other. These structural profiles are independent of variance of a particular structure (variance ratio analysis) or variance when considering multiple regions (Mahalanobis distance analysis). The cosine angle is a calculation of the dissimilarity between each individual’s structural profile and the centroid of their sex group. Larger average cosine angle indicates greater variance in the structural inter-relatedness profile of the respective sex.

The effects of age on cosine angle are notable for their developmental increase (and age-related decline). All of volume, surface area, and cortical thickness showed significant effects of age on cosine angle in the child and youth dataset. Cortical thickness notably showed the most prominent effects of age both in the developmental and late-life datasets. We interpreted these findings as a ‘biological validation’ of this method, in that differentiation of regions in cortical architecture has been shown to increase during neurodevelopment, and such differentiation decreases in late life with age-related change. The novel aspect that we have demonstrated here is that across an entire group, variability across individuals in relationships among regions increases during development, and (in cortical thickness) decreases in late life. When total brain volume was accounted for, age effects were stronger and more widespread across datasets and metrics (see supplement for full results).

#### Global Volume

Cosine angle was significantly different between females and males in the child and youth PNC dataset for global volume (females > males). There was a significant effect of age in both the PNC and the young adult HCP samples. There was also an age-by-sex interaction with global volume cosine angle in the OASIS-3 late-life dataset, such that male cosine angle was greater than female cosine angle in the latest-life group within this sample (Figure 4).

**Figure 4.**
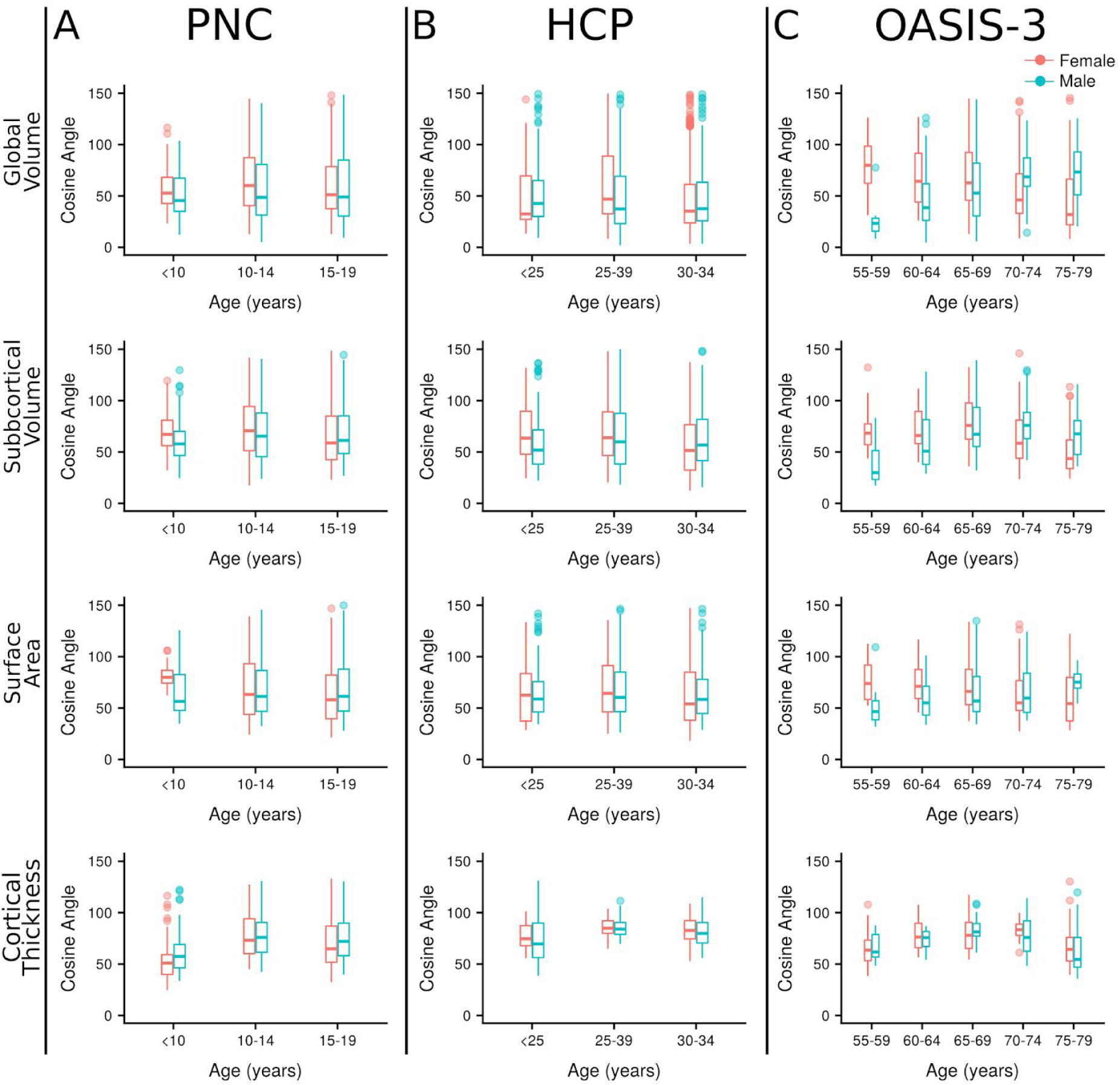
Cosine Angle Dissimilarity. Cosine angles are plotted by age bin for subjects from three independent datasets; (A) The Philadelphia Neurodevelopmental Cohort (PNC), (B) The Human Connectome Project (HCP) and (C) The Open Access Series of Imaging Studies (OASIS-3). Cosine angle was calculated for each subject to their group (male or female) centroid within each age bin separately on the surface of an n-dimensional sphere. Metrics included in each analysis were grouped by type; global volumes (n=4), subcortical volumes (n=14), surface area (n=68) and cortical thickness (n=68).

#### Subcortical Volume

No main effects of sex were present in cosine analysis of subcortical volume. There were significant effects of age in each sample, and a significant age-by-sex interaction in each sample. Specifically females had a greater cosine angle in early life, but males had a greater cosine angle in late life.

#### Surface Area

Within the surface area analyses there were no significant effects of sex. There was a significant effect of age in the child and youth PNC dataset. There was also an age-by-sex interaction in children and youth, where females displayed a greater cosine angle in early life (<10 years) while at older ages males and females were similar. In late life (OASIS-3) a similar age-by-sex interaction occurred, where males had a greater cosine angle compared to females in older groups of this late-life dataset.

#### Cortical Thickness

Significant effects of sex were present in cortical thickness in the child and youth PNC and young adult HCP datasets. In youth, cosine angle was greater in males compared to females. In the young adult sample it was greater in females compared to males. No age-by-sex interactions were present in the cortical thickness analysis. Age was significantly associated with cortical thickness cosine angle across all datasets. In all samples an inverted-U shape profile with age was seen with cosine angle first increasing with age before declining.

## Discussion

Across three large datasets and more than 3,000 MRI scans, we examined brain structural variability in global volume, and by region in subcortical volumes, surface area, and cortical thickness. We found that males consistently demonstrate greater variability in global brain volumes, subcortical volumes, and most notably surface area compared to females across the lifespan. This finding was stable across age. We then confirmed these univariate findings using the multivariate Mahalanobis distance approach, which showed that subcortical volumes and surface area were more variable in males compared to females across the lifespan. We then used the cosine angle metric to examine sex-based variability in how brain regions relate to each other. Only cortical thickness and global volume showed a main effect of sex, and this was early in the lifespan. Main effects of age were more prominent early and late in the lifespan across measures. Additionally, there were significant age-by-sex interactions at both ends of the lifespan for global volume, subcortical volume and surface area. When taken together, our results show that regional variability is greater in males compared to females. It starts early in the lifespan, and remains relatively constant. In contrast, variability among regions differs by sex in early verses late life. The greater regional variability in males in aspects of brain structure largely determined early with a strong genetic component, is consistent with the greater risk for males in neurodevelopmental disorders such as ASD and ADHD, which themselves have high heritability. Higher variability among brain regions, which may reflect greater specialization or maturation (indirectly supported by main effects of age, particularly during development), is greater in early life in females compared to males, but in the latest life group greater in males compared to females. These findings are consistent with greater neurodevelopmental disorder risk in boys, and greater dementia risk in women.

From the first report of sex differences in brain morphology in the early 1980’s (10), the topic has been controversial (11) and the evidence for sex differences in brain structure, not attributable to differences in brain size, is limited (15). The theory of sex differences in brain variability, rather than larger or smaller brain size, originated as far back as Darwin (33) and is supported by evidence across various species and phenotypes (34), primarily demonstrating a higher degree of variability in males. It is also unclear to what extent socio-political opinions have influenced decisions regarding publication of studies related to variability differences (e.g. (35, 36)). Examining variability in brain structure is an important endeavor as it may generate insights into the etiology of brain disorders, which often demonstrate differences in prevalance and disease course by sex. Furthermore, variability as a brain phenotype in its own right has become a topic of great interest in both healthy and brain disorder research groups (15–17).

Here we started by corroborating previous findings showing greater variability in male cortical surface area and subcortical regions in adults (44-77 years) (15) and extend this finding to data including participants with age ranging from childhood to late life. We also corroborate another former study showing greater male variability in subcortical volumes across development (3-21 years) (17). We built on these findings by using a lifespan approach across three datasets to show that males are more variable than females in subcortical volume and cortical surface area, a pattern which starts early and is sustained through late life. Additionally, as in Ritchie et al (15), we show that these variability differences are independent of variability in total brain volume. We then used a different approach to confirm our findings through calculation of the Mahalanobis distance for each brain phenotype. Using Mahalanobis distance allowed us to quantify the total magnitude of each individuals’ dissimilarity to their group average without assuming independence of regions to each other. We speculate that the age-by-sex interactions with surface area in the aging population may relate to the rate of atrophy differing between the sexes. It has been reported previously that the rate of volume loss is higher in males compared to females (37–39). Unlike surface area, we found similar regional variability of cortical thickness in both sexes across the lifespan. Independent genetic factors drive cortical thickness and surface area development (40–42), and these properties of brain structure demonstrate divergent developmental trajectories (43). Additionally, surface area developmental trajectories have been found to be sexually dimorphic, while trajectories of cortical thickness are similar for males and females (43). Cortical thickness is also thought to vary more in relation to environmental effects than surface area (41).

We speculate that mechanisms involved in the early propagation of intermediate radial glia cells, which are involved in the tangential expansion of the cortex (44), may be related to the variability differences seen in surface area between the sexes. The animal literature provides hints as to how these sex differences may arise as it has been shown that the estrogen steroid hormone estradiol, which is the major female sex hormone, promotes the proliferation of progenitor cells during development (45). Additionally, there is evidence that hormones modulate epigenetic regulation during development (46). Hill (35) posits a selectivity bias resulting in greater male variability while others have suggested greater variability is a consequence of the sex chromosomes architecture, where the heterogametic (i.e. XY in human males) structure leads to higher variability (47). The presence of two X chromosomes may play a protective role and create a blueprint for slightly less variability in brain structure in females compared to males. Recent work on sex-chromosome aneuploidy has indicated a dose effect of sex-chromosomes on gene expression (48). Additional work from the same group has found effects of sex-chromosome dosage on brain structures (49, 50). These studies show sex-chromosomes influence both brain structure and gene-expression, however, further work is required to identify the underlying causes of the discrepancy in variability between males and females seen here and determine the relationship to genetic variability or gene expression.

Although not identified by GWAS (40, 41), there is one X-chromosome gene that can influence cortical surface area; methyl-CpG binding protein 2 (MeCP2). Mutation of this gene, which is located at Xq28, is best known as the cause of Rett Syndrome, which almost exclusively affects females (51). In line with the theory of the protective X-chromosome, males with similar mutations suffer severe neonatal encephalopathy which is normally lethal within the first year of life (52). However, not all MeCp2 variations have such severe consequences. A common variant, rs2239464, has been shown in two independent datasets to be associated with reduced cortical surface area in males only (53). The effect was not seen in females. This was proposed to be due to the presence of a second copy of the gene in females protecting them (53). MeCP2 is a gene expression regulator (activator or repressor) of thousands of other genes (54) and potentially a key player influencing sex differences in surface area varibility shown here. Furthermore, animal work has shown interactions between sex hormones, MeCP2 and epigenetic regulation (46), however further research is required to fully elucidate these.

Our novel approach of investigating variability between regions using cosine angle analyses revealed differences in brain structure by sex that we do not believe has been previously reported. We observed significant age-by-sex interactions across all three datasets in the variability of structural profiles of subcortical volumes. Additionally, age-by-sex interactions were found during development and during later life in surface area. These interactions in children and youth may relate to differential rates of development (and regional differentiation), which are ‘slower’ in males compared to females (55). This aligns with previous reports of sexually dimorphic trajectories of development, at least for surface area (43), and the notion that different regions are under semi-independent genetic control (29, 40, 41, 56). In the latest life groups of older adults (OASIS-3), variability in the relationship among regions (in global volume and surface area) was greater in males than females. We can speculate that our findings relate to a consistent pattern of atrophy across the female population drawing them closer to their group mean. In contrast, findings in males could be due to atrophy occurring in more variable regions or more variable age of onset of atrophy across the population resulting in increased variability in structural profiles with increasing age (global volume and surface area). It is possible that greater variability among regions may be a protective mechanism against brain disorders, consistent with the finding of greater variability in early life in girls compared to boys, and in late life in males compared to females. This divergent relationship between the sexes with age in surface area, global and subcortical volumes might act to increase vulnerability for brain disorders in late life such as dementia (37), where the prevalence of Alzheimer’s disease (7) is increased in females compared to males.

Significant age effects were present in cortical thickness cosine angle across all datasets. Network approaches, using either structural covariance (grey matter volume and cortical thickness) or diffusion MRI, have shown significant age related changes in network architecture (30–32), characterized by network reorganization in aging. A separate study of structural covariance of cortical thickness highlighted the similarity in network architecture between males and females in terms of small-world organization, efficiency and node vulnerability where no sex differences were seen (57). These studies are in line with the current finding of an association between cortical thickness cosine angle and age, where cosine angle may reflect variability in the occurrence of these organizational changes across the population or the increased effect of environmental influences on cortical thickness.

This study has multiple major strengths, including the use of three large high quality datasets which encompass individuals from childhood to old age. Additionally, multiple complementary and advanced statistical approaches were applied to further understand healthy brain structural variation and elucidate variability both across and within individuals. Moreover, the use of these datasets and approaches allowed the replication of findings across samples and across methods providing strong evidence that they are generalizable. However, there are also some limitations to consider. There was an age difference between males and females in all three datasets. All analyses were repeated with age matched groups to ensure this was not skewing findings (supplementary material). The Desikan-Killiany atlas is somewhat limited in regional specificity, but was chosen to minimise the number of tests required and to make analyses comparable with previous studies. Furthermore, the majority of findings in the current study are so widespread (surface area for example) that the parcellation used was more than sufficient to detect differences. Finally, all data included in the current study that largely covered the lifespan were cross-sectional. It would be advantageous, although challenging, in future to utilise longitudinal data across the lifespan.

In conclusion, we demonstrated that males are more variable compared to females in individual brain regions with regard to surface area and volume measures. However, this increased variability in males does not extend to how brain regions relate to each other. Structural profile analyses highlighted a different aspect of variability demonstrating that the relationships between regions vary in a sex, age and metric specific fashion. In the future it would be of interest to investigate the association between sex differences in variability and genetic factors. For instance, how variability relates to gene expression maps may reveal fundamental information about driversity of these variability differences and potentially help uncover genetic risk factors for specific psychiatric disorders. Additionally, applying similar analyses to clinical populations could provide evidence of how neurostructural variability can result in vulnerability to psychiatric disease.

## Materials and methods

### Datasets

Basic demographic data are shown in Table 5.

**Table 4.** Cosine Angle Dissimilarity.

|  |  | PNC |  | HCP |  | OASIS-3 |  |
| --- | --- | --- | --- | --- | --- | --- | --- |
|  |  | F | q | F | q | F | q |
| Global | Sex | 6.83 | **0.03** | 1.25 | 0.26 | 1.40 | 0.26 |
|  | Age | 4.84 | **0.01** | 6.14 | **0.007** | 1.13 | 0.34 |
|  | Age-by-sex | 2.48 | 0.08 | 3.66 | **0.03** | 13.18 | **3.4E-10** |
| Subcortical Volume | Sex | 1.41 | 0.71 | 0.49 | 0.73 | 0.01 | 0.94 |
|  | Age | 7.09 | **0.001** | 6.38 | **0.002** | 9.88 | **3.3E-07** |
|  | Age-by-sex | 4.09 | **0.02** | 3.64 | **0.03** | 10.94 | **1.7E-08** |
| Surface Area | Sex | 0.06 | 0.81 | 0.06 | 0.81 | 1.32 | 0.75 |
|  | Age | 6.30 | **0.006** | 2.97 | 0.08 | 0.98 | 0.42 |
|  | Age-by-sex | 7.88 | **4.0E-04** | 0.85 | 0.43 | 7.33 | **9.8E-06** |
| Cortical Thickness | Sex | 9.16 | **0.008** | 6.61 | **0.02** | 1.63 | 0.20 |
|  | Age | 55.92 | **1.6E-23** | 42.47 | **2.8E-18** | 21.38 | **3.0E-16** |
|  | Age-by-sex | 2.66 | 0.07 | 1.18 | 0.31 | 1.55 | 0.19 |

Cosine angle distributions for three independent datasets; Philadelphia Neurodevelopmental Cohort (PNC), Human Connectome Project (HCP) and Open Access Series of Imaging Studies (OASIS-3) were compared between males and females with a linear model followed by type 2 F-tests. *q* - False Discovery Rate (FDR) corrected *p*-value. Cosine angles were calculated for each subject per age bin to their group (male or female) centroid on the *n*-dimensional sphere per metric type. Males and females were matched on age before analysis.

**Table 5.** Demographics.

|  | PNC (N=1,347) |  | HCP (N=1,032) |  | OASIS-3 (N=609) |  |
| --- | --- | --- | --- | --- | --- | --- |
|  | male | female | male | female | male | female |
| n | 615 | 732 | 507 | 606 | 240 | 369 |
| Age, mean (SD)* | 14.14 (3.48) | 14.61 (3.50) | 27.90 (3.61) | 29.56 (3.60) | 68.51 (8.97) | 66.54 (9.10) |
| Age, range | 8-21 | 8-21 | 22-36 | 22-37 | 42-89 | 43-95 |

\*There were significant age differences between males and females in each cohort. HCP - Human Connectome Project, OASIS-3 - Open Access Series of Imaging Studies, PNC - Philadelphia Neurodevelopmental Cohort, SD - standard deviation.

#### Child and Youth - PNC

Data for participants (n=1,601, aged 8-23) were included from the publicly available PNC dataset (21). Participants were scanned on the same 3T Siemens TIM Trio scanner at the Hospital of the University of Pennsylvania. T1-weighted images were acquired with a magnetization prepared, rapid-acquisition gradient-echo (MPRAGE) sequence with the following parameters: TR=1810 ms, TE=3.5 ms, TI=1100 ms, 9° flip angle and matrix of 192 × 256, resulting in a resolution of 0.94 × 0.94 × 1 mm^3^. Subjects were excluded based on missing data/processing errors (n=122), quality control (see Quality Control section below, n=51) and the presence of a major medical condition (n=81; e.g. epilepsy, skull fracture, meningitis, multiple sclerosis) resulting in n=1,347 individuals for analysis. On average females were slightly older than their male counterparts (Kruskal-Wallis = 6.88, *p*=0.01).

#### Young Adult - HCP

The HCP Young Adult S1200 (age 22-37) data were used (24). Only broadly healthy individuals were recruited to participate. T1-weighted data were collected on a custom 3T Siemens Skyra with a 3D MPRAGE sequence with the following parameters: TR=2400 ms, TE=2.14 ms, TI=1000, 8° flip angle and matrix of 320 × 256, resulting in a 0.7 mm^3^ isotropic resolution (24). High quality processed data were available for n=1,113 subjects. Females were older than males on average (Kruskal-Wallis = 73.46, *p*<0.0001).

#### Late-Life - OASIS-3

Data from cognitively normal aging adults (n=609, 43-95 years) from the OASIS-3 database were used (26–28). All MRI data included in this analysis were collected on one of two Siemens TIM Trio 3T MRI scanners at the Knight Alzheimer’s Disease Research Center, Washington University in St Louis with the following T1-weighted parameters: TR=400 ms, TE=3.16 ms, TI=1000 ms, 8° flip angle and matrix of 256 × 256, resulting in a 1 mm^3^ isotropic resolution. Due to the availability of longitudinal data, all 609 cognitively healthy subjects had data of good quality to include from a period when cognitively normal. Males were slightly older than females (Kruskal-Wallis = 7.51, *p*=0.006).

### Data Processing

FreeSurfer was used to segment subcortical structures and generate tessellated, smoothed grey-white and pial surfaces from the T1-weighted data (58–61). PNC data were processed in-house with FreeSurfer (v6.0). Metrics were then extracted from regions of the Desikan-Killiany parcellation (62), which includes 34 cortical and 7 sub-cortical regions per hemisphere. Similar summary metrics were directly available for download for the HCP and OASIS-3 datasets. HCP and OASIS-3 data used here were processed with FreeSurfer version 5.2 (enhanced version) (63) and 5.3, respectively.

Volume for global (total brain volume [TBV; BrainSeg_No_Vent or the sum of total cerebral and cerebellar grey and white matter], cerebral and cerebellar grey and white matter) and the 7 subcortical structures (thalamus, caudate, putamen, pallidum, nucleus accumbens, hippocampus and amygdala) were investigated along with cortical thickness and surface area from the 34 cortical regions per hemisphere.

### Quality Control (QC)

PNC FreeSurfer outputs were visually inspected in-house to ensure image quality and accurate segmentation of the grey and white matter. HCP data were quality controlled before being released. Similarly, the OASIS-3 FreeSurfer data were visually checked before release (64) - only data that passed inspection were included.

### Statistical analysis

All statistical analyses and graph generation were completed in the R statistical environment version 3.4.3 (65).

### Variance Ratio Across Measures and Regions

The variance associated with age was regressed from our measures leaving residuals that were then used for analyses. A generalized additive model (GAM, mgcv package) was applied to allow age to be modelled nonlinearly, avoiding the assumption of a linear, quadratic or cubic association between age and the metrics. The residuals were then Z-scored to provide measures of a similar magnitude for comparison. To compare variance between males and females, a variance ratio (VR) was generated with an F test (var.test). Next, to investigate the role of total brain volume in sex differences similar variance tests were conducted on Z-scored residuals from GAM models where total brain volume (linear) in addition to age (smooth) were regressed out. A linear term for total brain volume was deemed appropriate based on (66).

### Mahalanobis Distance

To refine our analyses, we grouped our metrics (corrected for age and Z-scored) by type; global volumes (n=4), subcortical volumes (n=14), surface area (n=68) and cortical thickness (n=68) and calculated Mahalanobis distance for each subject to their group average. For each metric type all measures of that type were included per subject. Mahalanobis distance is calculated as the distance from each subject to their group centroid - a multi-dimensional centre point representing the ‘average’ male and ‘average’ female set of metrics - while also accounting for the covariance of metrics. The metric used to account for the correlation structure utilised a covariance matrix computed over the full sample including both males and females. Thus a higher group average Mahalanobis distance would indicate a greater dispersion of the data in a group relative to its centroid. This was tested with a Welch two sample *t*-test.

### Cosine Angle Dissimilarity

For the second aim, we considered the relationship of regions to each other within a subject. Similar to the Mahalanobis distance analysis metrics were grouped by type (metrics corrected for age and Z-scored). Thus we characterised the unique pattern of structural metrics, a structural profile, for each individual as a direction vector in multivariate space. This was done separately for each metric type. Two individuals with similar structural profiles would have similar angles from the origin in multivariate space. Using cosine angle, we calculated the similarity between each individual and their group centroid, per metric type. The group centroid was calculated by minimizing the total sum of geodesic distances from the centroid to every other point on a unit sphere using an iterative algorithm generalised to n-dimensional spheres (67, 68). Once the centroid was calculated, we computed the average angular deviation from the group centroid for each sex. Differences in angular deviation (cosine angle) between the sexes were tested with a *t*-test.

#### Age

Additionally to test the influence of age, Mahalanobis distances and cosine angles were regenerated from the data in which the age effect was not regressed out (data still Z-scored). Data were binned according to age. Within each bin, Mahalanobis distance and cosine angle were calculated between each subject and their sexes centroid for that measure. Binning the data ensured that the age distribution or differences in the age distributions between males and females did not influence findings. To ensure adequate numbers were present in each bin the OASIS-3 sample was limited to 55-80 years. Linear models with type 2 F-tests were then used to examine the age-by-sex bin interaction.

False discovery rate (FDR) correction was implemented to account for multiple testing (*q*<0.05) within each analysis (i.e. univariate global analyses were corrected for n=5, subcortical analysis were corrected for n=14, surface area and cortical thickness analyses were corrected for n=68; multivariate analyses were corrected by 4, for the number of metric types).

Figure 5 provides a simplified 3-dimensional example of the two multivariate approaches.

**Figure 5.**
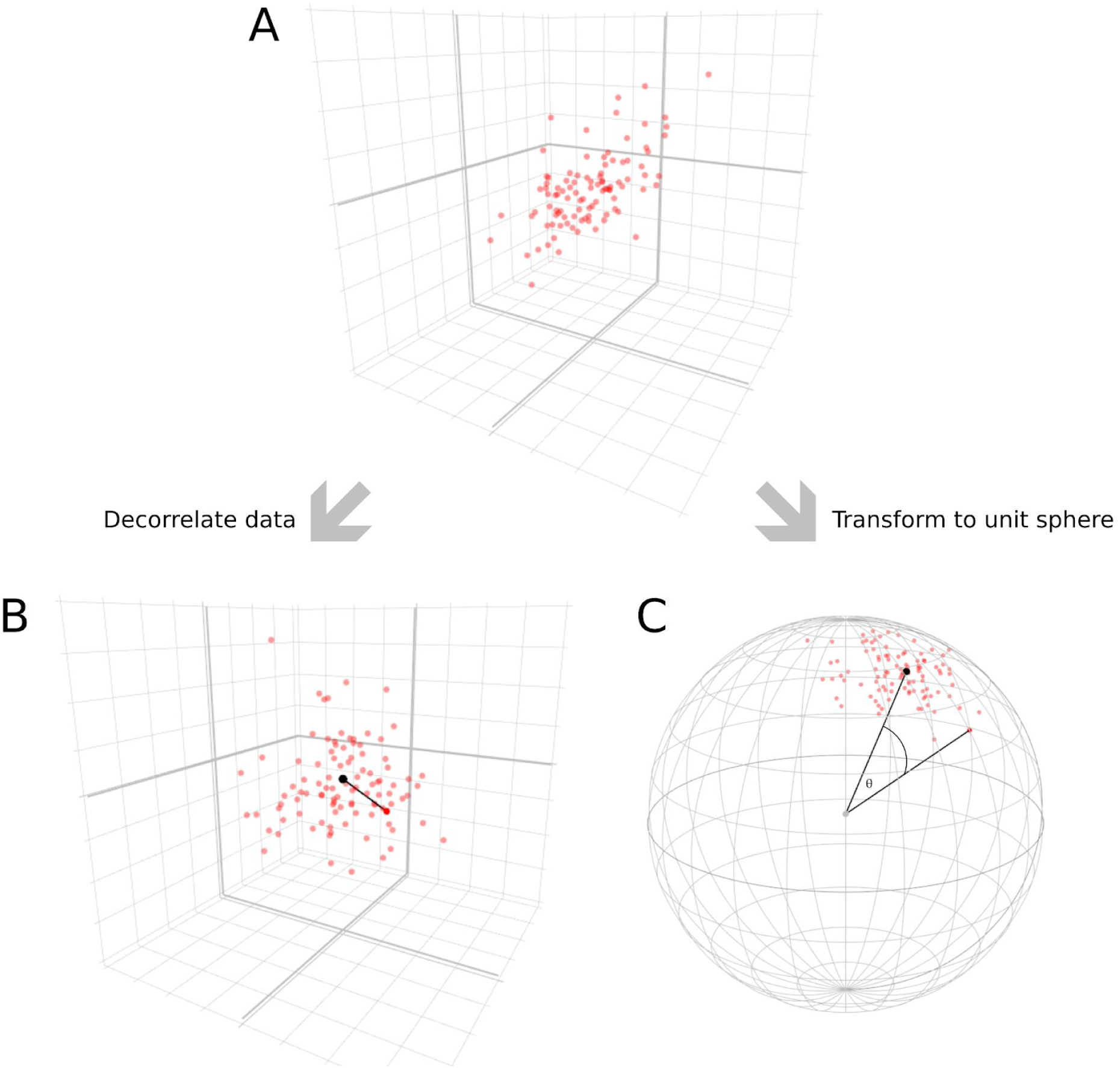
Simplified 3-dimensional schematic of the analytic methods. Each red point represents a subject per figure. (A) Raw data showing correlation between measures. (B) Displays data that has undergone a whitening or decorrelation step that occurs as part of Mahalanobis distance calculation. On decorrelated data the Mahalanobis distance is equivalent to the Euclidean distance between points. Distance (e.g. black line) is calculated between each subject (red point) and the data mean (black point) to give a multivariate measure of deviation based on distribution (or variance). (C) Shows data that has been normalised to the unit sphere. The magnitude of the measures are thus no longer represented but the proximity of points on the surface of the sphere indicates the correlational similarity of the subjects across all measures. The dissimilarity was quantified by calculating the cosine angle (*θ*) between each subject (red points) and the data centroid (black point). Black lines show an example.

## Acknowledgements

The Philadelphia Neurodevelopmental Cohort (PNC) was funded through NIMH RC2 grants MH089983 (Raquel E Gur) and MH089924 (Hakon Hakonarson).

Data were provided [in part] by the Human Connectome Project, WU-Minn Consortium (Principal Investigators: David Van Essen and Kamil Ugurbil; 1U54MH091657) funded by the 16 NIH Institutes and Centers that support the NIH Blueprint for Neuroscience Research; and by the McDonnell Center for Systems Neuroscience at Washington University

Data were provided [in part] by OASIS; OASIS-3: Principal Investigators: T. Benzinger, D. Marcus, J. Morris; NIH P50AG00561, P30NS09857781, P01AG026276, P01AG003991, R01AG043434, UL1TR000448, R01EB009352. OASIS grants: P50 AG05681, P01 AG03991, R01 AG021910, P50 MH071616, U24 RR021382, R01 MH56584.

