## supplemental information for "Sex differences in Variability of Brain Structure Across the Lifespan"

### Supplementary Material

#### Mahalanobis Distance

#### S Table 1 Mahalanobis Distance - T-tests - Age regressed out (TBV not regressed)

| | PNC | | | HCP | | | OASIS-3 | | |
|---|---|---|---|---|---|---|---|---|---|
| | t | q | d | t | q | d | t | q | d |
| Global Volume | -2.53 | **0.02** | -0.14 | -6.39 | **1.1E-09** | -0.42 | -4.64 | **9.0E-06** | -0.43 |
| Subcortical Volume | -3.96 | **1.1E-04** | -0.22 | -6.90 | **3.6E-11** | -0.44 | -6.86 | **4.6E-11** | -0.64 |
| Surface Area | -12.99 | **7.1E-36** | -0.75 | -15.78 | **1.4E-49** | -1.01 | -10.30 | **1.9E-22** | -0.95 |
| Cortical Thickness | -1.29 | 0.26 | -0.07 | 1.35 | 0.26 | 0.08 | 2.07 | 0.15 | 0.18 |

#### S Table 2 Mahalanobis distance - linear model - (TBV regressed out)

| | | PNC | | HCP | | OASIS-3 | |
|---|---|---|---|---|---|---|---|
| | | F | q | F | q | F | q |
| Global | Sex | 5.30 | **0.03** | 17.67 | **8.6E-05** | 1.60 | 0.21 |
| | Age | 0.29 | 0.75 | 1.02 | 0.54 | 4.92 | **0.002** |
| | Age-by-sex | 0.05 | 0.95 | 0.12 | 0.89 | 0.40 | 0.81 |
| Subcortical Volume | Sex | 16.32 | **5.7E-05** | 41.10 | **6.6E-10** | 39.86 | **9.3E-10** |
| | Age | 0.06 | 0.94 | 0.72 | 0.94 | 0.37 | 0.94 |
| | Age-by-sex | 0.56 | 0.57 | 0.76 | 0.47 | 2.42 | **0.047** |
| Surface Area | Sex | 159.07 | **2.8E-34** | 234.65 | **2.0E-47** | 21.75 | **4.1E-06** |
| | Age | 0.46 | 0.63 | 3.53 | **0.04** | 3.26 | **0.04** |
| | Age-by-sex | 2.23 | 0.11 | 1.43 | 0.24 | 3.59 | **0.007** |
| Cortical Thickness | Sex | 0.93 | 0.34 | 1.24 | 0.34 | 1.50 | 0.34 |
| | Age | 0.02 | 0.98 | 0.03 | 0.98 | 4.29 | **0.006** |
| | Age-by-sex | 0.57 | 0.57 | 0.09 | 0.92 | 3.21 | **0.01** |

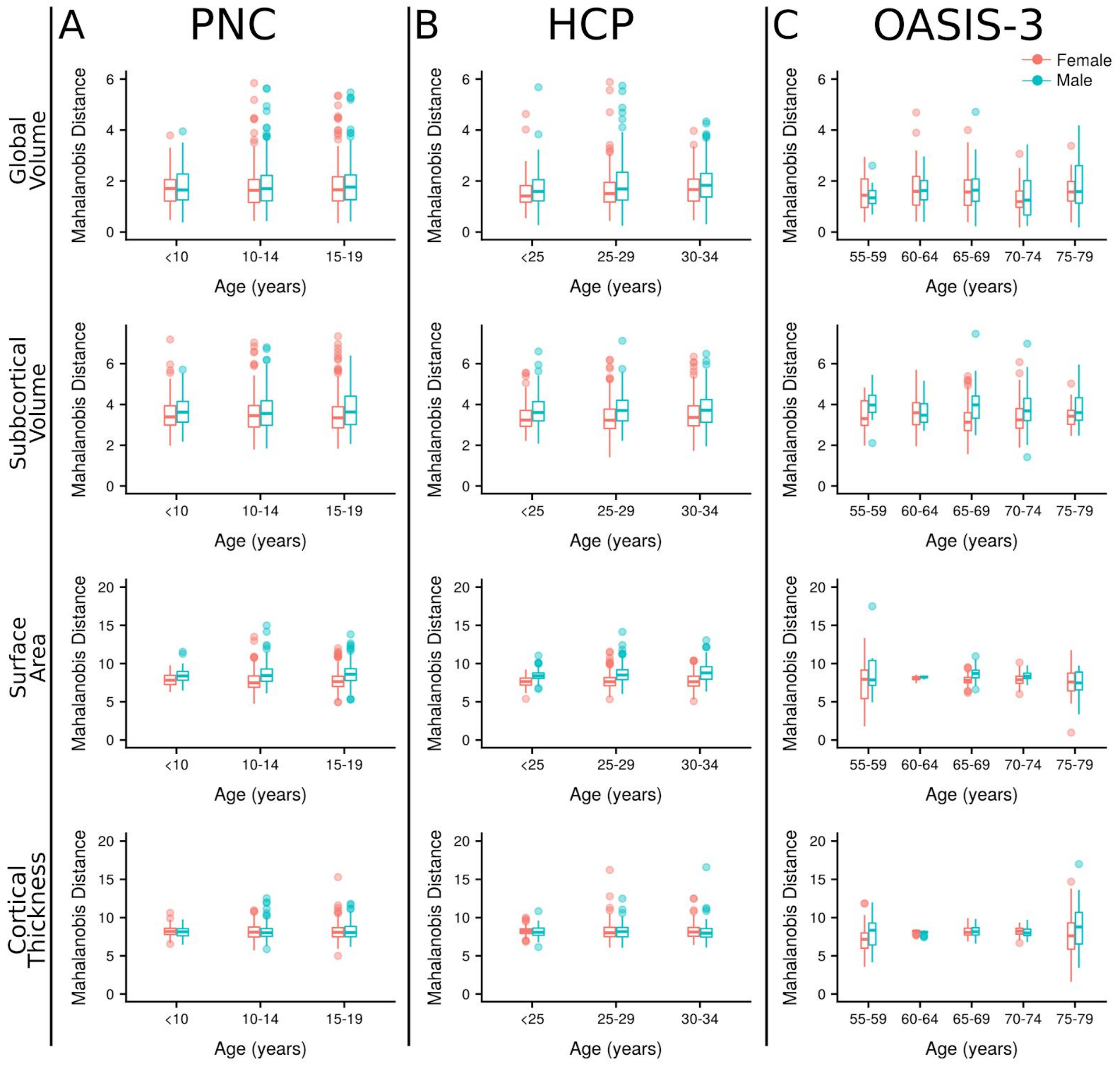


**S Figure 1 Mahalanobis distance (TBV regressed out)**

### Cosine dissimilarity

#### S Table 3 Cosine dissimilarity - t-test - Age regressed out (TBV not regressed)

| | PNC | | | HCP | | | OASIS-3 | | |
|---|---|---|---|---|---|---|---|---|---|
| | t | q | d | t | q | d | t | q | d |
| Global Volume | 1.31 | 0.35 | 0.07 | 0.13 | 0.90 | 0.01 | 1.20 | 0.35 | 0.11 |
| Subcortical Volume | 0.08 | 0.93 | 0.004 | -0.33 | 0.93 | -0.02 | 0.59 | 0.93 | 0.05 |
| Surface Area | -1.00 | 0.37 | -0.05 | -0.89 | 0.37 | -0.05 | 1.22 | 0.37 | 0.11 |
| Cortical Thickness | 1.33 | 0.18 | 0.07 | 1.39 | 0.18 | 0.09 | 1.33 | 0.18 | 0.12 |

#### S Table 4 Cosine dissimilarity - linear model - (TBV regressed out)

| | | PNC | | HCP | | OASIS-3 | |
|---|---|---|---|---|---|---|---|
| | | F | q | F | q | F | q |
| Global | Sex | 0.21 | 0.65 | 4.78 | **0.04** | 8.92 | **0.009** |
| | Age | 54.44 | **6.1E-23** | 50.37 | **2.0E-21** | 2.45 | 0.05 |
| | Age-by-sex | 6.37 | **0.002** | 1.08 | 0.34 | 1.51 | 0.20 |
| Subcortical Volume | Sex | 2.50 | 0.26 | 0.13 | 0.72 | 1.88 | 0.26 |
| | Age | 55.37 | **2.6E-23** | 8.40 | **2.4E-04** | 9.86 | **1.7E-07** |
| | Age-by-sex | 3.74 | **0.02** | 0.28 | 0.75 | 3.72 | **0.005** |
| Surface Area | Sex | 0.36 | 0.75 | 0.10 | 0.75 | 6.12 | **0.04** |
| | Age | 42.51 | **4.0E-18** | 14.27 | **7.7E-07** | 10.64 | **4.3E-08** |
| | Age-by-sex | 4.77 | **0.009** | 7.38 | **0.001** | 2.91 | **0.02** |
| Cortical Thickness | Sex | 4.66 | 0.05 | 5.26 | 0.05 | 2.42 | 0.12 |
| | Age | 52.65 | **3.2E-22** | 25.75 | **1.8E-11** | 10.06 | **7.9E-08** |
| | Age-by-sex | 5.93 | **0.003** | 8.93 | **1.4E-04** | 5.45 | **2.7E-04** |

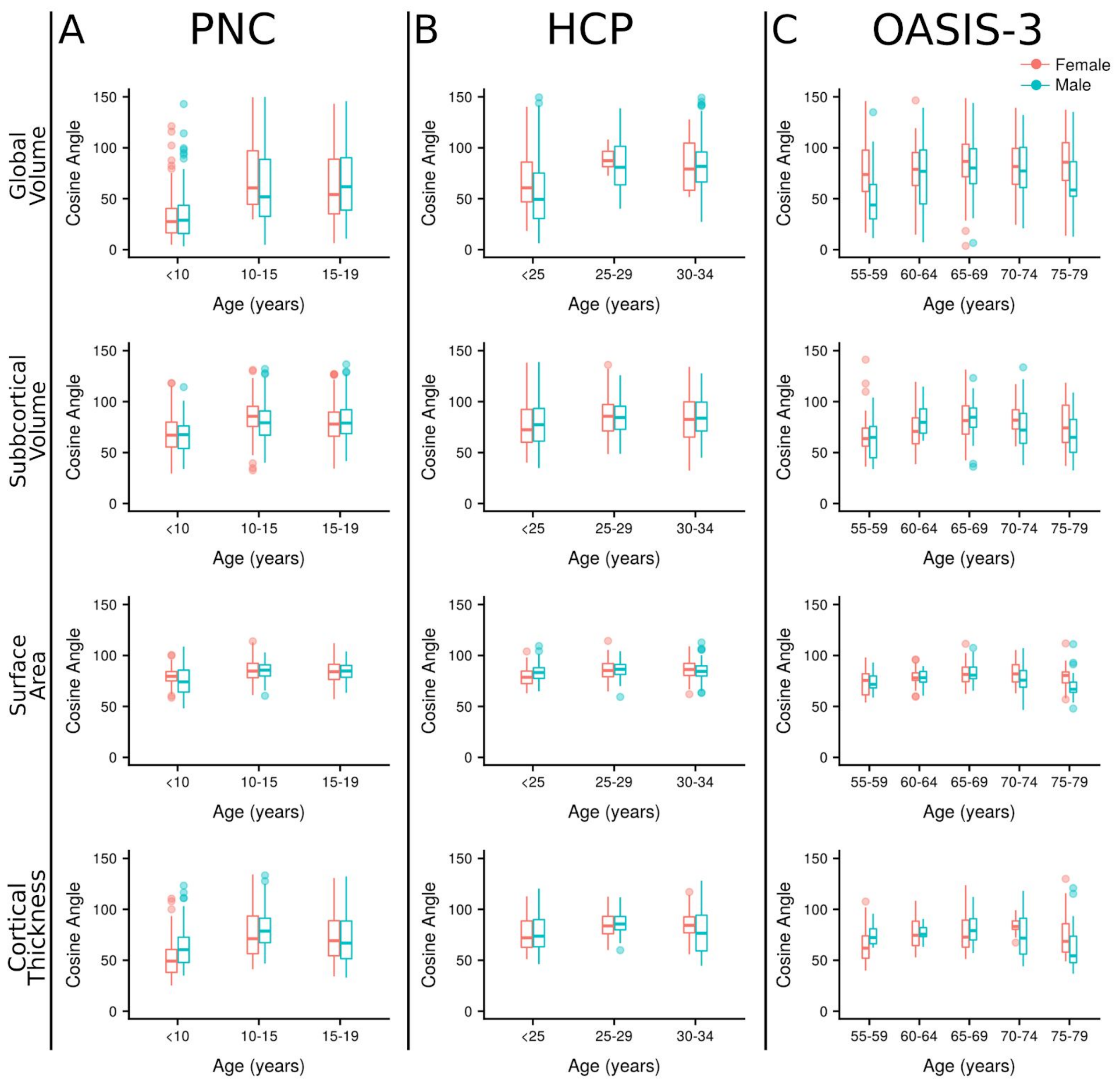


**S Figure 2 Cosine dissimilarity (TBV regressed out)**

#### Matching and Verification

Matching samples based on age (MatchIt, propensity scoring) had minimal impact on VR analysis (data not shown), therefore results from the full samples are reported in the main manuscript. For both multivariate analyses age bins were used which ensures differences in the overall age distributions would not affect findings.

**Mean sex effects**

*T*-tests were conducted to examine the difference on average between males and females. To evaluate the effect size of these differences we calculated Cohen's d (T statistic * 2 / √degrees of freedom).

*Volume*

Global and subcortical volumes were consistently higher in males compared to females on average, with medium to large effect sizes across all datasets (S Table 5&6, S Figure 3). Accounting for TBV rendered the majority of these non-significant, however cerebellar grey matter (all datasets), bilateral pallidum (PNC), left amygdala (PNC and HCP) and right amygdala (HCP) volumes remaining significantly larger in males compared to females. In contrast, the right hippocampus in PNC was larger in females compared to males after TBV correction (Cohen's d = 0.15).

| | | **PNC** | | | **HCP** | | | **OASIS-3** | | |
|---|---|---|---|---|---|---|---|---|---|---|
| | | t | q | d | t | q | d | t | q | d |
| TBV | raw | -23.29 | **2.5E-99** | -1.31 | -25.59 | **7.1E-111** | -1.62 | -24.66 | **1.1E-102** | -1.63 |
| Cerebral GM | raw | -22.71 | **2.6E-95** | -1.28 | -25.23 | **1.1E-108** | -1.58 | -24.18 | **8.4E-100** | -1.59 |
| | corrected | -1.61 | 0.25 | -0.09 | -1.77 | 0.13 | -0.11 | -1.44 | 0.19 | -0.10 |
| Cerebral WM | raw | -20.74 | **6.3E-82** | -1.18 | -22.88 | **5.7E-93** | -1.45 | -22.18 | **6.7E-87** | -1.48 |
| | corrected | 1.12 | 0.33 | 0.06 | 2.01 | 0.11 | 0.13 | 1.72 | 0.15 | 0.11 |
| Cerebellar GM | raw | -14.31 | **6.1E-43** | -0.80 | -21.62 | **9.6E-86** | -1.34 | -21.07 | **2.7E-81** | -1.35 |
| | corrected | -3.24 | **0.01** | -0.18 | -5.53 | **2.0E-07** | -0.34 | -5.48 | **2.8E-07** | -0.35 |
| Cerebellar WM | raw | -4.86 | **1.7E-06** | -0.27 | -16.37 | **2.6E-53** | -1.05 | -16.22 | **4.2E-52** | -1.07 |
| | corrected | 1.43 | 0.25 | 0.08 | -1.42 | 0.20 | -0.09 | -1.68 | 0.15 | -0.11 |

**S Table 5** Mean differences in global volumes

Results from analyses of sex differences in global brain volumes are presented from the Philadelphia Neurodevelopmental Cohort (PNC), Human Connectome Project (HCP) and Open Access Series of Imaging Studies (OASIS-3). Mean differences in male and female volumes were compared with age regressed out (raw) and age and total brain volume (TBV) regressed out (corrected). Mean differences were compared with a *t*-test (t). Positive Cohen's d (d) represents females > males while negative represents males > females on average.

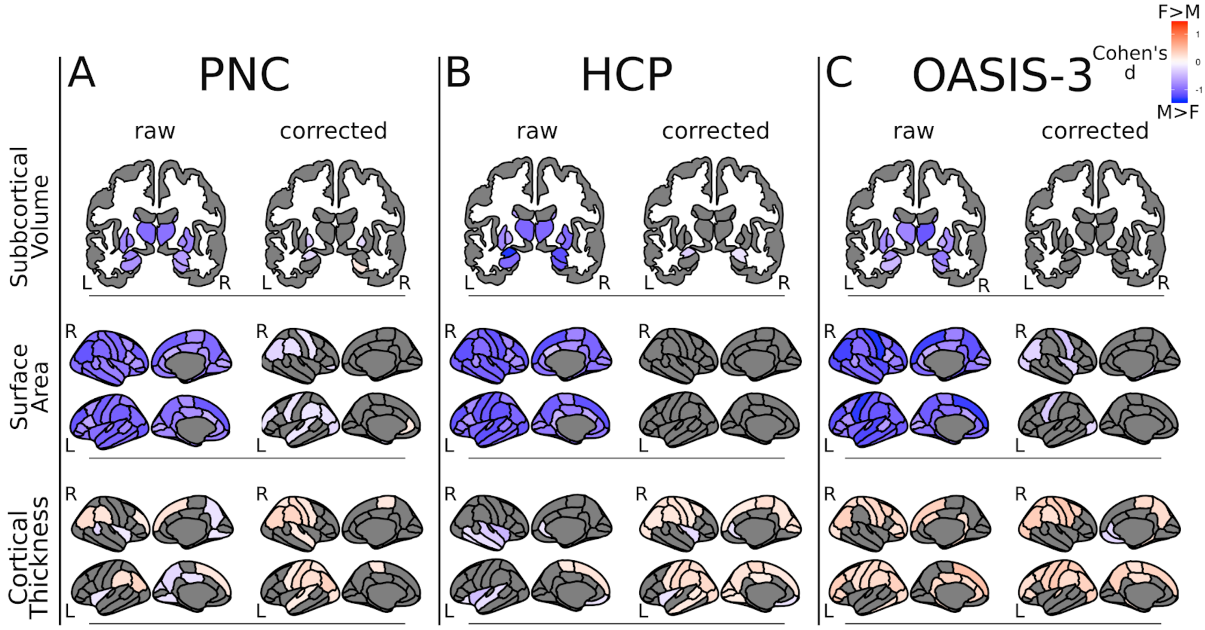


**S Figure 3** Average sex differences
Cohen's d effect sizes are mapped onto subcortical structures (top row) or cortical surface (middle and bottom panels) for three independent datasets; (A) The Philadelphia Neurodevelopmental Cohort (PNC), (B) The Human Connectome Project (HCP) and (C) The Open Access Series of Imaging Studies (OASIS-3). These results were generated with *t*-tests to compare average differences in subcortical volume, cortical surface area and cortical thickness between males and females. Only Cohen's d from tests that met statistical significance are plotted ($q<0.05$). Cohen's d > 0 (red) indicates females > males, Cohen's d < 0 (blue) indicates males (M) > females (F). 'Raw' figures show results from analyses that used metrics corrected for age (age effects regressed out). Corresponding 'corrected' figures show results from analyses that used metrics corrected for total brain volume (TBV) as well as age. Note: Nucleus Accumbens is not included in the figure but does show significant volume differences on average between males and females (males larger) across all samples in the raw analysis (see S Table 6). L- left, R - right.

| | | | PNC | | | HCP | | | OASIS-3 | | |
|---|---|---|---|---|---|---|---|---|---|---|---|
| | | | t | q | d | t | q | d | t | q | d |
| Thalamus | left | raw | -17.67 | **1.7E-61** | -0.99 | -16.48 | **7.5E-54** | -1.03 | -8.96 | **3.4E-17** | -0.82 |
| | | corrected | -1.07 | 0.45 | -0.06 | 1.78 | 0.15 | 0.11 | -0.09 | 0.93 | -0.01 |
| | right | raw | -16.87 | **8.7E-57** | -0.96 | -16.26 | **1.4E-52** | -1.03 | -11.45 | **2.5E-25** | -1.14 |
| | | corrected | 0.10 | 0.92 | 0.01 | 1.25 | 0.28 | 0.08 | -0.65 | 0.88 | -0.06 |
| Caudate | left | raw | -11.16 | **1.4E-27** | -0.63 | -10.79 | **9.2E-26** | -0.67 | -6.18 | **2.1E-09** | -0.61 |
| | | corrected | 0.53 | 0.69 | 0.03 | 2.10 | 0.08 | 0.13 | 0.15 | 0.93 | 0.01 |
| | right | raw | -11.59 | **2.2E-29** | -0.65 | -11.80 | **3.9E-30** | -0.74 | -7.31 | **2.6E-12** | -0.71 |
| | | corrected | -1.22 | 0.39 | -0.07 | 1.54 | 0.19 | 0.10 | -0.58 | 0.88 | -0.06 |
| Putamen | left | raw | -14.12 | **8.7E-42** | -0.80 | -13.08 | **3.5E-36** | -0.80 | -7.88 | **7.5E-14** | -0.76 |
| | | corrected | -1.66 | 0.23 | -0.09 | -1.60 | 0.19 | -0.10 | -0.85 | 0.79 | -0.08 |
| | right | raw | -14.95 | **5.9E-46** | -0.84 | -16.55 | **3.0E-54** | -1.02 | -7.54 | **7.4E-13** | -0.75 |
| | | corrected | -1.55 | 0.24 | -0.09 | -2.21 | 0.08 | -0.14 | -0.99 | 0.79 | -0.10 |
| Pallidum | left | raw | -14.83 | **1.8E-45** | -0.84 | -9.99 | **1.5E-22** | -0.61 | -8.00 | **3.8E-14** | -0.75 |
| | | corrected | -6.02 | **3.2E-08** | -0.34 | -2.15 | 0.08 | -0.13 | -2.78 | 0.08 | -0.26 |
| | right | raw | -13.56 | **6.9E-39** | -0.77 | -12.23 | **3.8E-32** | -0.75 | -5.95 | **7.0E-09** | -0.56 |
| | | corrected | -4.16 | **2.4E-04** | -0.24 | -0.46 | 0.76 | -0.03 | 0.14 | 0.93 | 0.01 |
| Hippocampus | left | raw | -13.19 | **3.7E-37** | -0.73 | -15.66 | **3.9E-49** | -1.01 | -5.03 | **7.6E-07** | -0.47 |
| | | corrected | 0.71 | 0.61 | 0.04 | -1.22 | 0.28 | -0.08 | 1.86 | 0.30 | 0.17 |
| | right | raw | -11.54 | **2.7E-29** | -0.64 | -17.88 | **6.9E-62** | -1.11 | -6.71 | **9.2E-11** | -0.64 |
| | | corrected | 2.77 | **0.02** | 0.15 | -2.16 | 0.08 | -0.13 | 1.35 | 0.62 | 0.13 |
| Amygdala | left | raw | -14.60 | **2.5E-44** | -0.82 | -21.62 | **3.7E-84** | -1.37 | -7.29 | **2.6E-12** | -0.70 |
| | | corrected | -3.90 | **4.7E-04** | -0.22 | -5.05 | **7.2E-06** | -0.32 | -0.15 | 0.93 | -0.01 |
| | right | raw | -11.55 | **2.7E-29** | -0.65 | -19.20 | **1.3E-69** | -1.20 | -9.40 | **2.1E-18** | -0.89 |
| | | corrected | -1.65 | 0.23 | -0.09 | -2.85 | **0.03** | -0.18 | -2.06 | 0.28 | -0.19 |
| Nucleus Accumbens | left | raw | -4.35 | **1.5E-05** | -0.24 | -13.11 | **3.1E-36** | -0.82 | -3.77 | **1.8E-04** | -0.35 |
| | | corrected | -0.99 | 0.45 | -0.06 | -0.33 | 0.77 | -0.02 | -0.92 | 0.79 | -0.09 |
| | right | raw | -8.54 | **4.0E-17** | -0.48 | -13.40 | **1.2E-37** | -0.84 | -5.48 | **8.4E-08** | -0.52 |
| | | corrected | -0.29 | 0.83 | -0.02 | -0.29 | 0.77 | -0.02 | -0.46 | 0.91 | -0.04 |

**S Table 6** Mean differences in subcortical volumes
Results from analyses of sex differences in subcortical brain volumes are presented from the Philadelphia Neurodevelopmental Cohort (PNC), Human Connectome Project (HCP) and Open Access Series of Imaging Studies (OASIS-3). Mean differences in male and female volumes were compared with age regressed out (raw) and age and total brain volume (TBV) regressed out (corrected). Mean differences were compared with a *t*-test (t). Positive Cohen's d (d) represents females > males while negative represents males > females on average.

*Surface area*

Similarly, SA was significantly higher in males compared to females across the cortex in all datasets, accounting for TBV again reduced these effects. However, some regions did remain significantly larger in males compared to females in the PNC (12 regions) and OASIS-3 samples (10 regions) datasets (S Figure 3). Following TBV correction there was also one region in the development sample (PNC) where SA was significantly greater in females; left rostral anterior cingulate (Cohen's d 0.16, $q<0.05$, S Figure 3).

*Cortical Thickness*

Results from the analysis of CT was more variable across datasets. Females had greater CT than males in various regions across all datasets with small to medium effect sizes. While males also displayed greater CT than females in certain regions (S Figure 3). Accounting for TBV reduced the extent of differences where males were greater than females and increased the number of regions where females had significantly larger CT than males.
